# Midbrain-Hippocampus Structural Connectivity Selectively Predicts Motivated Memory Encoding

**DOI:** 10.1101/2022.05.18.492387

**Authors:** Blake L. Elliott, Kimberlee D’Ardenne, Vishnu P. Murty, Gene A. Brewer, Samuel M. McClure

## Abstract

Motivation is a powerful driver of learning and memory. Functional MRI studies show that interactions between the dopaminergic midbrain (SN/VTA), hippocampus, and nucleus accumbens (NAc) are critical for motivated memory encoding. However, it is not known if these effects are transient and purely functional, or if individual differences in the structure of this circuit underlie motivated memory encoding. To quantify individual differences in structure, diffusion-weighted MRI and probabilistic tractography were used to quantify SN/VTA-striatum and SN/VTA-hippocampus pathways associated with motivated memory encoding in humans.

Participants completed a motivated source memory paradigm. During encoding, words were randomly assigned to one of three conditions: reward ($1.00), control ($0.00), or punishment (-$1.00). During retrieval, participants were asked to retrieve item and source information of the previously studied words and were rewarded or penalized according to their performance. Source memory for words assigned to both reward and punishment conditions was greater than control words, while there were no differences in item memory based on value. Anatomically, probabilistic tractography results revealed a heterogeneous, topological arrangement of the SN/VTA. Tract density measures of SN/VTA-hippocampus pathways were positively correlated with individual differences in reward and punishment modulated memory performance, while density of SN/VTA-striatum pathways showed no association. This novel finding suggests that pathways emerging from the human SV/VTA are anatomically separable and functionally heterogeneous. Individual differences in structural connectivity of the dopaminergic hippocampus-VTA loop are selectively associated with motivated memory encoding.

**Significance Statement:** Functional MRI studies show that interactions between the dopaminergic midbrain (SN/VTA), hippocampus, and nucleus accumbens (NAc) are critical for motivated memory encoding. This has led to competing theories that posit either SN/VTA-NAc reward prediction errors or SN/VTA-hippocampus signals underlie motivated memory encoding. Additionally, it is not known if these effects are transient and purely functional, or if individual differences in the structure of these circuits underlie motivated memory encoding. Using diffusion-weighted MRI and probabilistic tractography, we show that tract density measures of SN/VTA-hippocampus pathways are positively correlated with motivated memory performance, while density of SN/VTA-striatum pathways show no association. This finding suggests that anatomical individual differences of the dopaminergic hippocampus-VTA loop are selectively associated with motivated memory encoding.

---

The motivation to obtain rewards and avoid punishments is a powerful driver of learning and memory. Scientists have investigated the effect of reward motivation on human episodic memory for nearly a century (Heyer Jr & O’Kelly, 1949; Kahneman & Peavler, 1969; Loftus & Wickens, 1970; Weiner & Walker, 1966). In cognitive neuroscience, most studies have focused on functional properties of the dopamine system rather than more stable neuroanatomical differences. We used diffusion weighted imaging to investigate structural connectivity of neural circuits involved in reward- and punishment-motivated memory encoding.

The neural mechanisms underlying reward-motivated memory encoding remain unresolved. Modulatory effects of the dopaminergic reward system on memory-related brain regions of the medial temporal lobe are hypothesized to be critical (Knowlton & Castel, 2022; Shohamy & Adcock, 2010). Neuroimaging studies in humans have found the involvement of the midbrain substantia nigra/ventral tegmental area (SN/VTA) and the nucleus accumbens (NAc; the main component of the ventral, limbic striatum) in the prediction of both rewards and punishments (Carter et al., 2009; McClure et al., 2003; Samanez-Larkin et al., 2007). Results from studies implementing functional magnetic resonance imaging (fMRI) during reward-motivated memory encoding have shown that increased activation in the medial temporal lobe and the reward system (i.e. SN/VTA and NAc) are predictive of better memory prioritization for higher-valued stimuli (Adcock et al., 2006; Shigemune et al., 2014; Wittmann et al., 2005).

However, it is not clear if this pattern of activation represents canonical SN/VTA-NAc reward prediction errors observed during reinforcement learning (Calderon et al., 2021; Jang et al., 2019; Schultz, 1998), SN/VTA-hippocampus reward/novelty signals (Lisman et al., 2011; Lisman & Grace, 2005), or some combination of the two (Ergo et al., 2020). To test between these possibilities, the present study tested whether the structural connectivity of one or both pathways is associated with behavioral measures of motived memory encoding.

Anatomically, animal research has revealed that the midbrain SN/VTA is not a uniform structure. The SN/VTA has a distinct topological organization, with heterogeneous parallel pathways from dopamine nuclei that support separate cognitive processes. The medial regions (VTA, medial SN) innervate the ventral (limbic) striatum, while more lateral regions innervate the dorsal (executive and motor) striatum (Hedreen & Delong, 1991; Lynd-Balta & Haber, 1994; Selemon & Goldman□Rakic, 1990; Szabó, 1979). The SN/VTA-striatum connections form an upward-spiraling circuit that integrates signals from affective, executive control, and motor regions to produce goal-directed behavior (Haber, 2016). The SN/VTA also forms a circuit with the hippocampus that is responsible for encoding information into long-term memory (i.e. the hippocampus-VTA loop; Lisman & Grace, 2005). The upward arc of the hippocampus-VTA loop consists of direct dopaminergic projections to the hippocampus (Gasbarri et al., 1991; Gasbarri et al., 1997; Lewis et al., 2001; Swanson, 1982). These neurons release dopamine into the hippocampus, which promotes synaptic plasticity by enhancing long-term potentiation (Lisman et al., 2011; Lisman & Grace, 2005).

Although the relationship between the anatomical circuitry of the SN/VTA and its functions has been documented in rodents and nonhuman primates, research in human populations has been limited to functional activation and functional connectivity. We used diffusion weighted MRI and probabilistic tractography to assess structural connectivity of the SN/VTA to reward- and punishment-motivated memory encoding. We hypothesized that the SN/VTA would be anatomically and functionally heterogeneous. Anatomically, we aimed to resolve the topological heterogeneity of the SN/VTA non-invasively in a neurotypical human population. Medial regions (VTA) were hypothesized to have higher connectivity with ventral (limbic) regions of the striatum and with the hippocampus. More lateral regions of the SN/VTA were hypothesized to have higher connectivity with dorsal (executive and motor) regions of the striatum. Given the fMRI findings implicating the SN/VTA, limbic striatum, and hippocampus in reward-motivated memory encoding, we hypothesized that structural connectivity of one or both tracts would be associated with reward- and punishment-motivated memory encoding.

## Method

### Participants

Fifty participants (17 male, 33 female) were recruited via an online newsletter and flyers posted around the Arizona State University campus (mean age 20.36 years, SD = 1.94, min = 18.21 years, max = 28.17 years). Participants were told that they could be compensated up to $100 for an MRI study investigating memory and decision-making. Participants were all right-handed and were screened for neurological or psychiatric disorders. Two subjects were excluded from the DTI analysis. One subject was excluded because they were missing the reverse phase-encoding scan needed for FSL topup distortion correction. The other subject was excluded because they did not have a usable structural scan. Two additional subjects were excluded from the final analyses because they scored at or below chance level on their overall memory performance (i.e. hit rates), leaving a final sample size of 46 participants.

### Experimental Paradigm

The experimental task closely replicated Shigemune et al. (2014). The task consisted of three repetitions of an encoding phase followed by a retrieval phase. The stimuli consisted of 704 words drawn randomly from the Penn Electrophysiology of Encoding and Retrieval Study word pool (an edited version of the Kucera-Francis word pool, available at http://memory.psych.upenn.edu/; Lohnas & Kahana, 2013). The encoding phases consisted of 30 words each randomly assigned to one of three conditions: reward ($1.00), control ($0.00), or punishment (-$1.00), 10 of each value. The word and value pairs were shown concurrently for 2s on either the left or right side of the screen with an interstimulus interval randomly jittered between 3s and 5s. Participants were instructed to encode both the word (item) and location (source) of the word. The paradigm was presented using MRI compatible video goggles and the task was completed in the scanner.

The retrieval phases consisted of 45 words (30 from the most recent list and 15 new words, randomly intermixed) presented one at a time in the center of the screen without values. The words were presented for 2s with an interstimulus interval randomly jittered between 3s and 5s. Participants were asked to judge whether the word was previously presented on the left side of the screen during the study phase, the right side of the screen, or whether the word was a “new” word that had not been studied before. If the word was previously in the reward condition, the participants were told they would gain 50¢ for correctly recognizing the word but getting the location incorrect, and an additional 50¢ if both the word and location were correct. If the word was previously in the punishment condition, the participants were told they would avoid losing 50¢ for correctly recognizing the word but getting the location incorrect and avoid losing an additional 50¢ if both the word and location were correct. If the word was previously in the control condition the participant neither gained nor lost any money based on their memory performance. The participants were told they would lose 50¢ for incorrectly classifying “new” words as “old” (i.e. false alarms). Participants were paid the total amount of money earned in the task at the end of the session. Responses were made using a five-button optic response box.

### Behavioral Data Analysis

Items with correct source memory (item with source hits: IWS) and items that were correctly classified as old but with incorrect source memory (item only hits: IO) were aggregated into proportions of total correct responses in that category by dividing by total number of studied items. The experimental design is a 3×2 repeated measures ANOVA with the following factors: condition (reward, control, punishment) and memory judgment (IWS, IO). Following Shigemune et al. (2014), we expected a main effect of condition and memory judgment. Importantly, we also expected a significant interaction between the two factors. Reward and punishment motivation were hypothesized to selectively enhance associative memory and not item memory. Therefore, we also expected a polynomial trend analysis to show a quadratic effect for the interaction between the two factors (i.e. reward > control < punishment in the IWS condition only).

### MRI Acquisition, Preprocessing, and Analysis

#### Structural MRI

MRI scans were acquired on a 3.0 T GE Discovery MR750 using a Nova Medical 32-channel head coil (General Electric, Milwaukee, WI). For each participant, high-resolution, T1-weighted structural images were collected using a co-planar, single shot, interleaved, 3D magnetization-prepared rapid gradient-echo (BRAVO) in the sagittal plane with the following imaging parameters: TR = 7.236 ms, TE = 2.784 ms, FOV 23.0 cm, 12° flip angle, slice thickness 0.9 mm, slices per slab=192, voxel size 0.9 × 0.9 × 0.9 mm, matrix size 236 × 256 × 256. The scan took approximately six minutes to complete.

#### Diffusion-weighted MRI

Diffusion-weighted images were collected using 48 directions, with a total gradient diffusion sensitivity of b = 1000 s/mm^2^ with 6 repeats of the B0 (no diffusion weighting) image, resulting in a 4D volume with the following imaging parameters: TR = 7800 ms, TE = 60.4 ms, FOV 23.2 cm, slice thickness 2.0 mm, voxel size 0.91 × 0.91 × 2.0 mm, matrix size 256 × 256 × 80 × 54. The scans were acquired posterior-anterior. The scan took approximately seven minutes to complete. An additional B0 image was collected in the reverse phase encoding direction (anterior-posterior) for use with FSL topup EPI distortion correction.

The diffusion-weighted data analysis closely follows the procedures from our previous study in which we quantified structural connectivity of SN/VTA-striatum tracts in a clinical sample, establishing the reliability of this protocol (Elliott et al., 2021). The diffusion-weighted data were processed using FSL (www.fmrib.ox.ac.uk/fsl). After correcting for movement and eddy current artifacts, the diffusion parameters were calculated for each voxel using FSL bedpostX. Measures of tract strength were calculated using probabilistic tractography, which is based on a probabilistic Bayesian framework (Behrens et al., 2007; Behrens et al., 2003). Fiber tracking was conducted in parallel for each voxel within a predefined SN/VTA seed mask. We used 5,000 samples per voxel, a curvature threshold of 0.2, and a step length of 0.5 mm. Target areas in the striatum were defined using a connectivity-based segmentation atlas with subdivisions for sensorimotor, executive, and limbic regions; this atlas is freely available with the FSL software (http://fsl.fmrib.ox.ac.uk/fsl/fslwiki/Atlases/striatumconn). These striatal subdivisions also likely reflect separable pathways from midbrain dopaminergic regions. The homogeneity of dopamine release measured with positron emission tomography in response to the administration of amphetamine is significantly higher within these connectivity-based subdivisions of the striatum compared to within anatomical subdivisions (i.e., putamen, caudate, and nucleus accumbens; Tziortzi et al., 2014). For specificity and to stay true to the anatomy, we refer to these striatal pathways as SN/VTA-motor, SN/VTA-executive, and SN/VTA-limbic when discussing our results. Hippocampus masks were defined using the Harvard-Oxford subcortical atlas integrated within FSL (Figure 1). The MNI space target masks were normalized to each participant’s native space using the inverse of the spatial normalization parameters. To tailor the ROIs to each individual’s anatomy, we used individually segmented gray matter (GM) and fractional anisotropy (FA) images to mask the ROIs. Following Tziortzi et al. (2014), the lower threshold for the GM mask was set at 0.25, and the FA mask upper threshold was set at 0.40. The seed ROI for the SN/VTA was defined using a probabilistic atlas of human SN/VTA (Murty et al., 2014). We used a 50% probability threshold, and the mask was normalized to each participant’s native space using the inverse of the normalization parameters.

**Figure 1:**
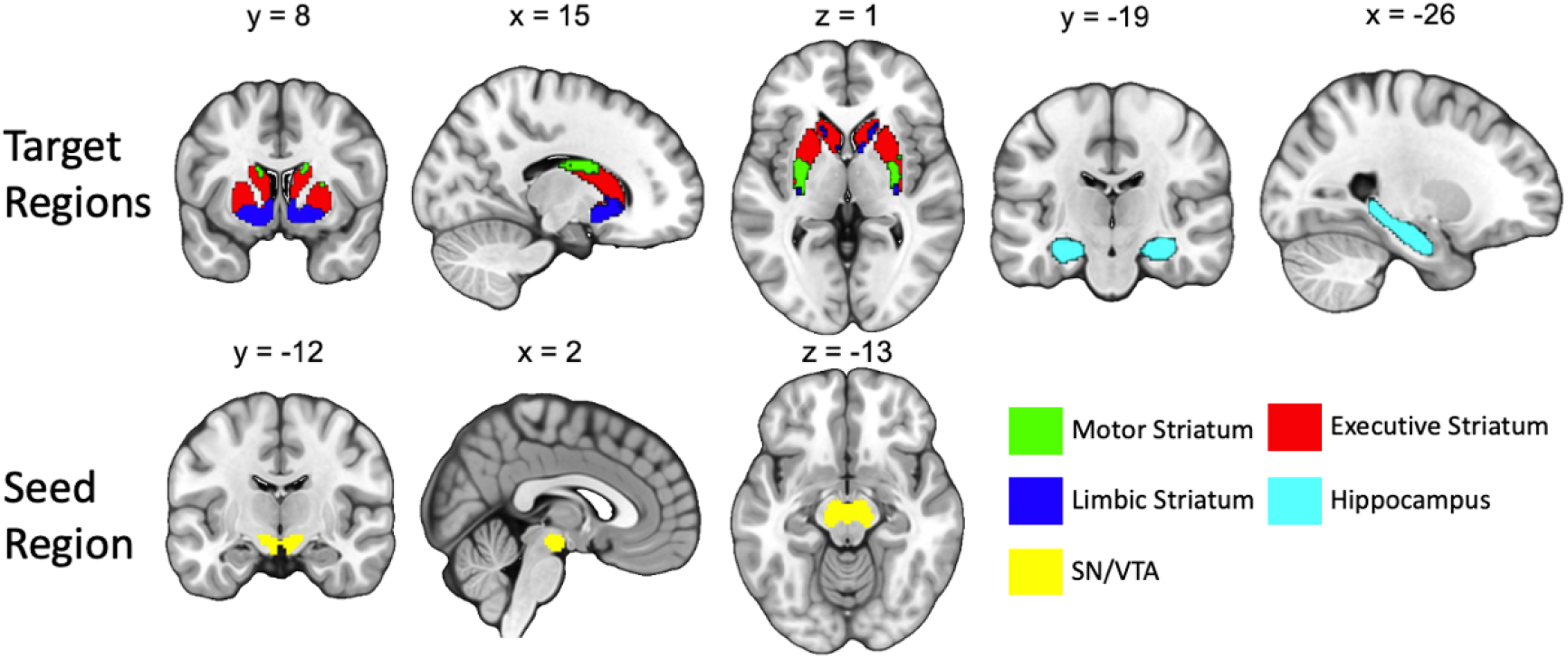
Seed and Target Regions for the probabilistic tractography analysis. The target regions of the striatum were functionally segmented based on projections from motor, executive, and limbic cortices (Tziortzi et al., 2014). The seed region of the midbrain was generated from a combined SN/VTA probabilistic atlas (Murty et al., 2014).

Because the dorsal and ventral tiers of the SN and VTA have not yet been reliably resolved in humans with MRI, we use the term “SN/VTA-striatum tracts” to describe meso- and nigro-striatal tracts. We note that the use of this term is not meant to imply directionality because diffusion-weighted imaging data do not include information about whether tracts are efferent or afferent. We chose to use the larger SN/VTA combined mask due to anatomical variability and the relatively small size of the SN and VTA (Halliday & Törk, 1986; Keuken et al., 2014). However, the results of the tractography analysis should follow our a priori predictions of the topology of the SN/VTA. Higher probability of connection with the hippocampus and limbic striatum should be localized in the medial SN and VTA. Higher probability of connection with the executive and motor striatum should be localized primarily in the SN.

All tractography analyses were conducted in the participants’ native anatomical space. The FSL FDT toolbox was used to perform probabilistic tractography with a partial volume model (Behrens et al., 2003), allowing for up to two fiber directions in each voxel (Behrens et al., 2007). We generated 5,000 sample tracts from each voxel in the SN/VTA seed mask. Visual inspection was used to ensure that the tractography maps were successful and acceptable for further analysis. Tractography was performed separately for the left and right striatum and hippocampus, and possible tracts were restricted to the hemisphere of origin using an exclusion mask of the contralateral hemisphere. Following standard procedures, the seed-based classification maps were first thresholded so that only voxels with at least 10 tracts terminating in one of the target regions were kept (Cohen et al., 2009; Forstmann et al., 2012). Next, the voxel values were converted into proportions of the number of tracts reaching the target mask from one voxel, divided by the number of tracts generated from that voxel (maximum 5,000). We used the mean of these value maps as the measure of SN/VTA-striatum and SN/VTA-hippocampus tract density. The term “tract integrity” is often used to describe microstructural measures (e.g. fractional anisotropy). To avoid confusion, we use the term “tract density”.

#### SN/VTA Topology

To determine the organization of SN/VTA-striatum and SN/VTA-hippocampus across participants, individual probability maps were normalized into Montreal Neurological Institute space. Each map was thresholded so that only voxels that had at least 50 samples reaching any target region were included (Tziortzi et al., 2014). The resulting maps were averaged to create 8 total for the entire sample (one for each target region for each hemisphere – limbic, executive, and motor striatum and hippocampus, Figure 4). Visual inspection and similarity metrics (Dice and dilated Dice coefficients) were used to quantify the topological arrangement of the probability maps in relation to the SN and VTA, respectively. Because the larger combined SN/VTA probability maps completely encapsulated both the SN and VTA, the maps were restricted to include only voxels that were greater than the mean value of the entire map. Thus, only voxels with the highest probability of connection were retained for comparison. Similarity metrics (Dice and dilated Dice coefficients) were calculated for the thresholded maps and the SN and VTA separately for each hemisphere (Table 1).

**Table 1.** Dice and dilated Dice statistics for the SN and VTA with each target region.

region
| Target Region | Dice |  | Dilated Dice |  |
| --- | --- | --- | --- | --- |
|  | SN | VTA | SN | VTA |
| Left Motor | 0.31 | 0.31 | 0.83 | 0.60 |
| Left Executive | 0.32 | 0.37 | 0.79 | 0.67 |
| Left Limbic | 0.23 | 0.52 | 0.67 | 0.79 |
| Left Hippocampus | 0.14 | 0.52 | 0.58 | 0.79 |
| Right Motor | 0.34 | 0.28 | 0.82 | 0.61 |
| Right Executive | 0.32 | 0.42 | 0.76 | 0.71 |
| Right Limbic | 0.30 | 0.49 | 0.71 | 0.80 |
| Right Hippocampus | 0.12 | 0.46 | 0.58 | 0.77 |

The Dice coefficient represents the degree of spatial overlap between two ROIs and was calculated as follows: 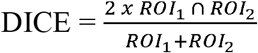. The dilated Dice coefficient is a suitable measure for small structures with complex shapes such as the VTA. This measure is similar to the Dice coefficient except that it dilates each mask by one voxel. This means that a single-voxel offset is not penalized as it is for the classical Dice coefficient, an important factor for small, complex structures. Both coefficients range from 0 to 1, with 0 representing no overlap and 1 representing complete overlap. Both measures were derived using the “segmentation_statistics” function from the Nighres python toolbox (Huntenburg et al., 2018). We expected that limbic and hippocampus maps would have higher Dice and dilated Dice values with the VTA and lower values with the SN. Conversely, the executive and motor maps were expected to have higher Dice and dilated Dice values with the SN and lower values with the VTA.

#### Tract Density and Motivated-Memory

The connectivity of each fiber tract was calculated by taking the mean of each probability map corresponding to each target region for each subject (four per hemisphere, eight in total). Hierarchical regressions were used to test the association of SN/VTA-hippocampus tracts and SN/VTA-limbic tracts with behavioral measures of reward and punishment prioritization (i.e. the difference between correct reward and control IWS hits, and the difference between correct punishment and control IWS hits). The diffusion measures for both the hippocampus and limbic tracts were averaged across the left and right hemispheres in order to circumvent the issue of multicollinearity in our regression models. Intracranial volume was entered in the first step of the model. The second step of the model included SN/VTA-hippocampus tracts and SN/VTA-limbic tracts, with intracranial volume treated as a covariate.

We further tested the association of SN/VTA-hippocampus tracts and SN/VTA-limbic tracts on reward and punishment prioritization separately for each hemisphere. Tract density measures were correlated across subjects with behavioral measures of reward and punishment prioritization. Partial correlations were conducted, with intracranial volume included as a covariate. Based on previous fMRI evidence implicating the hippocampus-VTA loop in reward-motivated memory encoding, we expected positive correlations between SN/VTA-hippocampus tract density and individual differences in reward- and punishment-motived encoding. The SN/VTA and the limbic striatum have also been implicated in reward-motivated memory, possibly due to reward prediction error signaling (Ergo et al., 2020). We therefore also conducted correlations between SN/VTA-limbic tract density and individual differences in reward- and punishment-motived encoding.

## Results

### Behavioral Results

Memory performance as a function of value and response type (correct item with source, IWS, and item only, IO, performance) is summarized in Figure 2. The results replicate Shigemune et al. (2014). Specifically, a 3 (condition: reward, control, punishment) × 2 (memory judgment: IWS, IO) repeated measures ANOVA with Greenhouse-Geisser corrections revealed a marginal effect of condition (*F*_*(1*.*427,47)*_ = 3.44, *p* = 0.053, η_p_^2^ = .07), and a main effect of memory judgment (*F*_*(1,47)*_ = 152.8, *p* < 0.001, η_p_^2^ = .77). We also found a significant interaction between the two factors (*F*_*(1*.*92,47)*_ = 4.24, *p* < 0.05, η_p_^2^ = .08). A quadratic trend analysis revealed a significant interaction of value and response type (*F*_*(1,47)*_ = 6.78, *p* < 0.05, η_p_^2^ = .13). These results demonstrate that both reward and punishment motivated encoding conditions increased memory for items that were accurately remembered with the correct source location (Figure 2).

**Figure 2:**
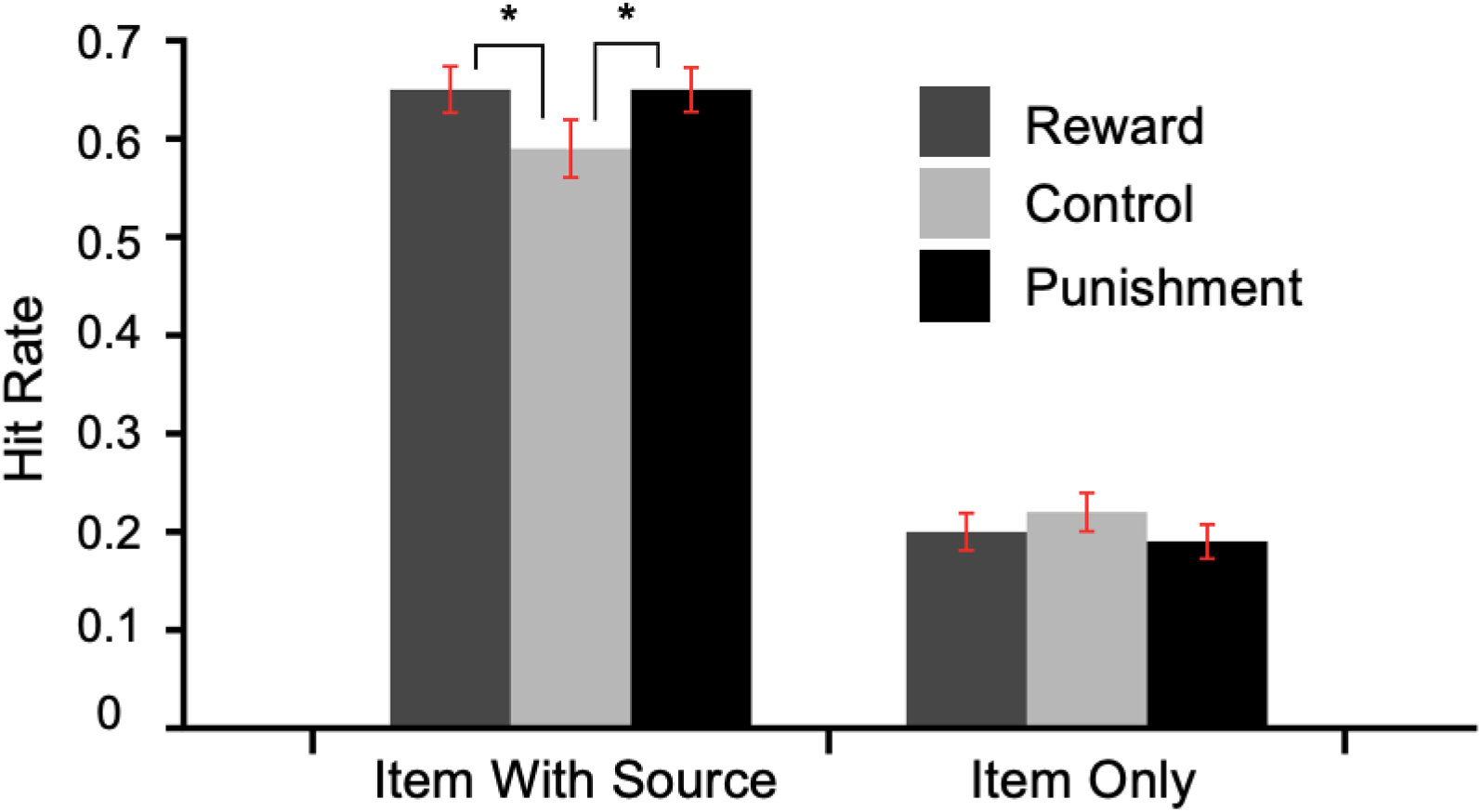
Memory performance as a function of encoding condition. IWS = item with source memory, IO = item only memory. Error bars represent 95% CI. * *p* < .05

We quantified individual differences in memory prioritization for reward and punishment as the difference between reward and control IWS hits and the difference between punishment and control IWS hits (Figure 3). We chose to use only IWS hits because this is where the effect of value was isolated in previous experiments with similar tasks (Elliott et al., 2020; Shigemune et al., 2014). This method has also been shown to form a stable latent factor with reward-modulated “remember” response in a VDR remember-know paradigm (Elliott et al., 2020). A one sample t-test showed that both the reward (M = .06, SD = .17) and punishment (M = .07, SD = .17) effects were statistically greater than 0: t(47) = 2.36, p < 0.05, Cohen’s d = .341; t(47) = 2.795, p < 0.05, Cohen’s d = .403. These effects were also strongly correlated (*r*(47) = .723, *p* < .001). Due to concerns that these effects may be driven by a participant with large reward and punishment effects (greater than 3 SD from the mean in both categories, Mahalanobis distance = 1.9; Extended Data Figure 3-1), we performed bootstrapped one sample t-tests with 1,000 samples for reward (95% CI .014 to .109, p < .05) and punishment (95% CI .024 to .117, p < .05). We also performed one sample t-tests with this subject removed. Both the reward and punishment effects were statistically greater than 0: t(46) = 2.16, p < 0.05, Cohen’s d = .315; t(46) = 2.75, p < 0.01, Cohen’s d = .401, and the measures were still highly correlated (Figure 3; *r*(46) = .549, *p* < .001). We chose to remove this outlier from the following connectivity correlation analyses. For completeness and transparency, scatterplots and statistics with this participant included can be found in the Extended Data Figure 3-1 and 5-1.

**Figure 3:**
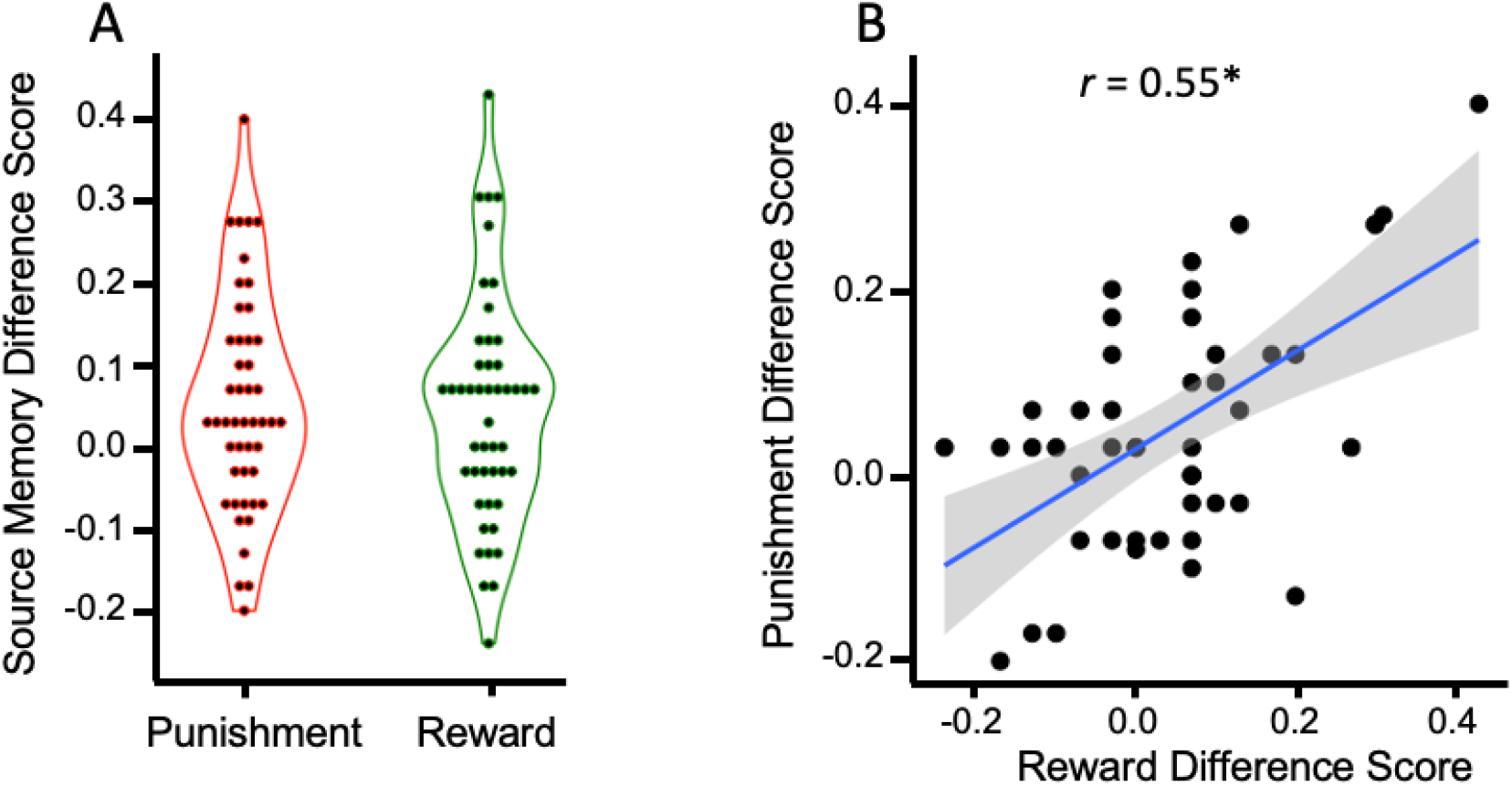
Reward and punishment prioritization effects. A) Violin plot showing reward (reward source memory hit rate minus control source memory hit rate) and punishment (punishment source memory hit rate minus control source memory hit rate) effects in source memory B) Scatterplot showing the significant correlation between reward and punishment source memory difference scores (* *p* < 0.001).

### Structural Organization of the SN/VTA

Before relating structural connectivity to motivated memory, we first wanted to confirm that the anatomical organization of the SN/VTA corroborates with what has been shown in rodents and non-human primates. When tracts with endpoints in the midbrain SN/VTA, functionally-parcellated striatal regions, and hippocampus were classified, four subdivisions emerged within the midbrain SN/VTA (Figure 4). The organization of these subdivisions was consistent with known anatomy in animals and non-human primates (Haber & Knutson, 2010). The subdivisions revealed an inverse dorsal-ventral and medial-lateral connectivity with the striatum. Projections from the SN/VTA to the limbic striatum were primarily located in the VTA and medial SN. Projections from the SN/VTA to the executive striatum were located more laterally, primarily in the SN. Projections from the SN/VTA the motor striatum were located most laterally in the SN. Additionally, connectivity between the SN/VTA and hippocampus was primarily localized within voxels located in the VTA.

**Figure 4:**
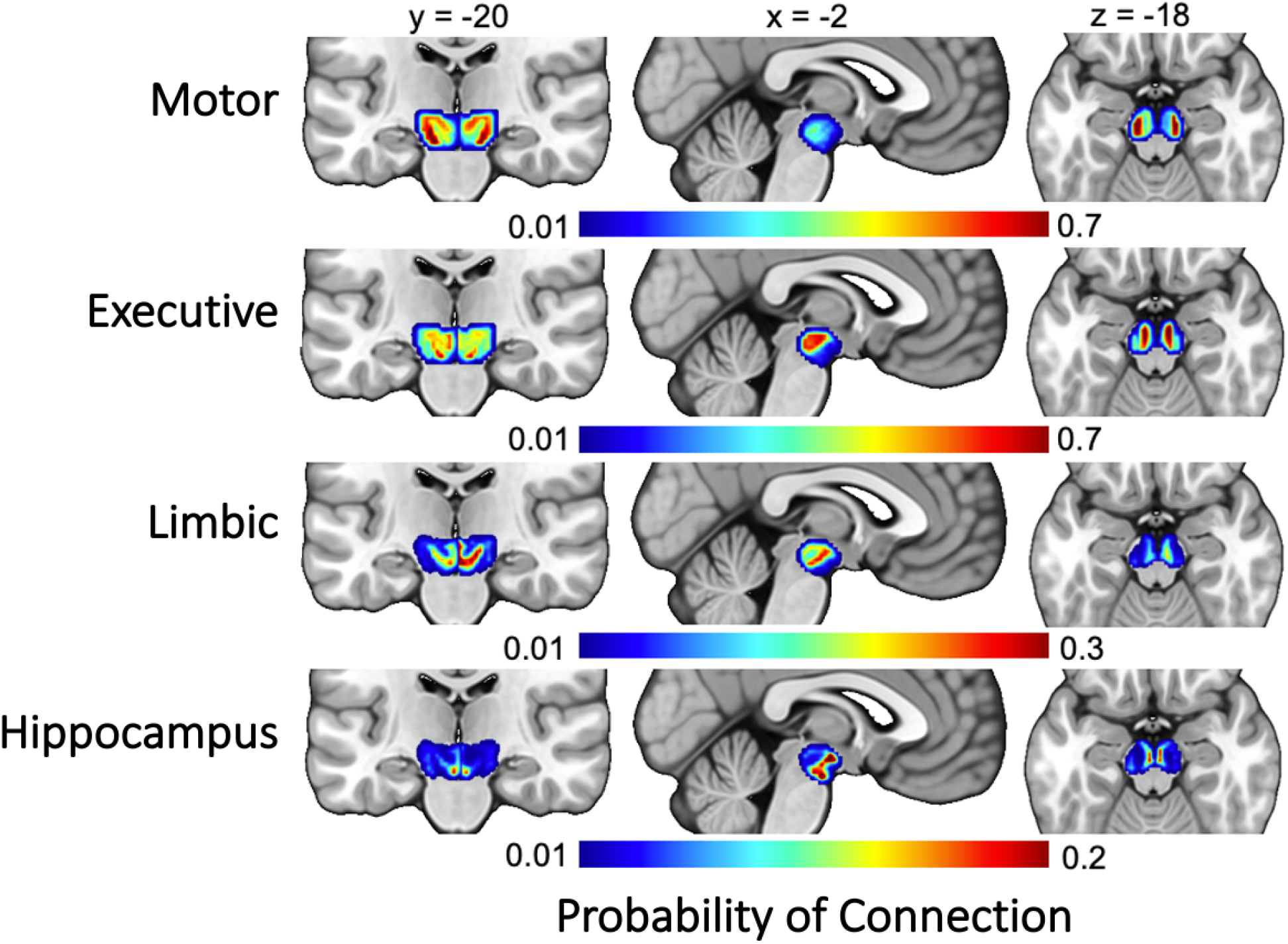
Group average projections from the SN/VTA to the four target regions (motor striatum, executive striatum, limbic striatum, hippocampus).

To further assess the topology of the SN/VTA projections, Dice and dilated Dice coefficients were calculated between each target region and the SN and VTA separately. Because the larger combined SN/VTA probability maps would completely encapsulate both the SN and VTA, these maps were restricted to include only voxels that were greater than the mean value of the entire map, leaving only the voxels with the highest probability of connection with each target region. The results are shown in Table 1. Both Dice and dilated Dice similarity metrics revealed a linear pattern of overlap with the SN and the target regions in both hemispheres.

Motor tracts had the largest Dice coefficients with the SN, followed by executive, limbic, and hippocampus. The opposite pattern was shown with the VTA: limbic and hippocampus tracts had the largest Dice coefficients, followed by executive and lastly, motor tracts. These findings comport with our a priori hypothesis and what is known from animal histology. We were able to demonstrate non-invasively in humans a topological arrangement of the midbrain SN/VTA.

### Tract Density and Motivated Memory Performance

We next conducted analyses to determine whether individual differences in reward-motivated memory prioritization were related to SN/VTA-hippocampus tract density using hierarchical multiple regression (Table 2). The first step of the hierarchical regression model included intracranial volume, so that the amount of variance explained by the structural connectivity measures can be isolated in the next step by examining the change in R^2^. Based on this first step, 10.7% of the variance in reward prioritization was accounted for by intracranial volume. Results from the second step revealed that SN/VTA-hippocampus tracts alone were associated with reward prioritization. Based on this additive model, 15.1% of the variance in reward prioritization could be accounted for by the structural connectivity measures (SN/VTA-hippocampus and SN/VTA-limbic tract density), controlling for intracranial volume (R square change = .151). Cohen’s *f*^2^ indicated that this was a medium effect (*f*^2^ = 0.18).

**Table 2.** Hierarchical linear regression models predicting reward prioritization.

| | B | SE | $\beta$ | *t* | *p* |
| --- | --- | --- | --- | --- | --- |
| **Step 1: Covariate** |  |  |  |  |  |
| Intracranial Volume | 1.05e<sup>-7</sup> | <0.001 | 0.107 | 0.703 | 0.486 |
| **Step 2: Additive Effect** |  |  |  |  |  |
| SN/VTA-Limbic | -0.071 | 0.504 | -0.021 | -0.140 | 0.889 |
| SN/VTA-Hippocampus* | 2.408 | 0.888 | 0.406 | 2.713 | 0.010 |

A similar hierarchical regression was conducted for punishment prioritization (Table 3). The first step of the hierarchical regression model included intracranial volume, so that the amount of variance explained by the structural connectivity measures can be isolated in the next step by examining the change in R^2^. Based on this first step, 0.7% of the variance in punishment prioritization was accounted for by intracranial volume. Results from the second step revealed that SN/VTA-hippocampus tracts alone were associated with punishment prioritization. Based on this additive model, 17.6% of the variance in reward prioritization could be accounted for by the structural connectivity measures (SN/VTA-hippocampus and SN/VTA-limbic tract density), controlling for intracranial volume (R square change = .176). Cohen’s *f*^2^ indicated that this was a medium effect (*f*^2^ = 0.21).

**Table 3.** Hierarchical linear regression models predicting punishment prioritization.

| | B | SE | $\beta$ | *t* | *p* |
| --- | --- | --- | --- | --- | --- |
| **Step 1: Covariate** |  |  |  |  |  |
| Intracranial Volume | 6.84e<sup>-9</sup> | <0.001 | 0.007 | 0.049 | 0.961 |
| **Step 2: Additive Effect** |  |  |  |  |  |
| SN/VTA-Limbic | -0.212 | 0.465 | -0.067 | -0.457 | 0.650 |
| SN/VTA-Hippocampus* | 2.422 | 0.818 | 0.439 | 2.960 | 0.005 |

We further tested the association of SN/VTA-hippocampus tracts and SN/VTA-limbic tracts on reward and punishment prioritization separately for each hemisphere. We computed partial correlations for the reward effect and SN/VTA-hippocampus tract density and the punishment effect and SN/VTA-hippocampus tract density for each hemisphere (controlling for intracranial volume). The results revealed that both the reward (*r*(42) = .315, *p* < .05) and punishment (*r*(42) = .325, *p* < .05) memory effects were positively correlated with the left SN/VTA-hippocampus tract. Both the reward (*r*(42) = .325, *p* < .05) and punishment (*r*(42) = .340, *p* < .05) were positively correlated with the right SN/VTA-hippocampus tract as well (Figure 5). We also conducted analyses to determine whether individual differences in memory prioritization were related to SN/VTA-limbic tract density. We computed partial correlations for the reward effect and SN/VTA-limbic tract density and the punishment effect and SN/VTA-limnic tract density for each hemisphere (controlling for intracranial volume). The results revealed that neither the reward (*r*(42) = -.159, *N*.*S*.) nor punishment (*r*(42) = -.122, *N*.*S*.) memory effects were correlated with the left SN/VTA-limbic tracts. Additionally, both the reward (*r*(42) = .221, *N*.*S*.) and punishment (*r*(42) = .112, *N*.*S*.) were not correlated with the right SN/VTA-limbic tracts as well (Figure 6).

**Figure 5:**
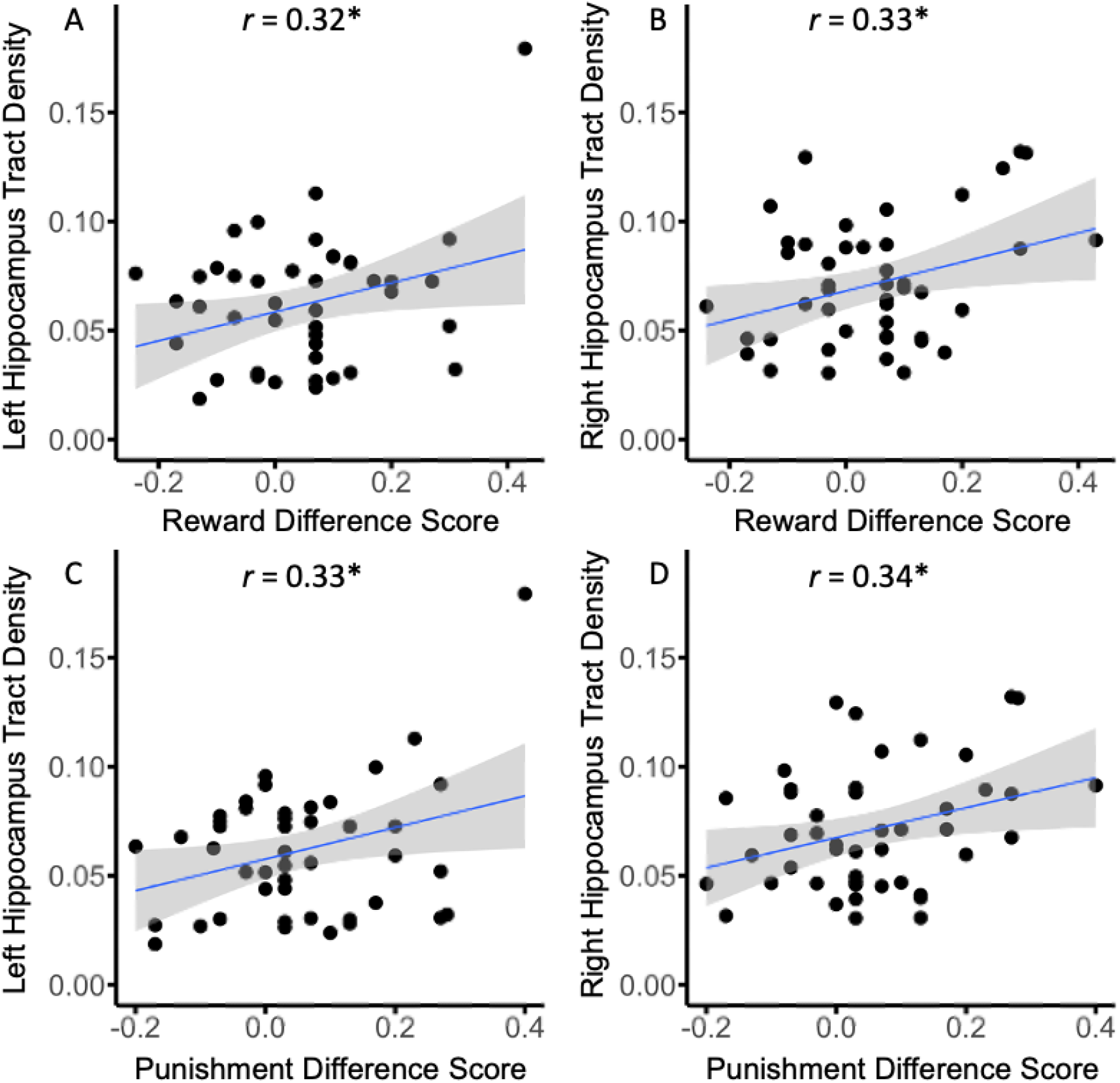
Scatterplots of reward (reward source memory hit rate minus control source memory hit rate) and punishment (punishment source memory hit rate minus control source memory hit rate) effects and hippocampus tract density. A) Left hippocampus tract density and reward effect B) Right hippocampus tract density and reward effect C) Left hippocampus tract density and punishment effect D) Right hippocampus tract density and punishment effect (* *p* < 0.05).

**Figure 6:**
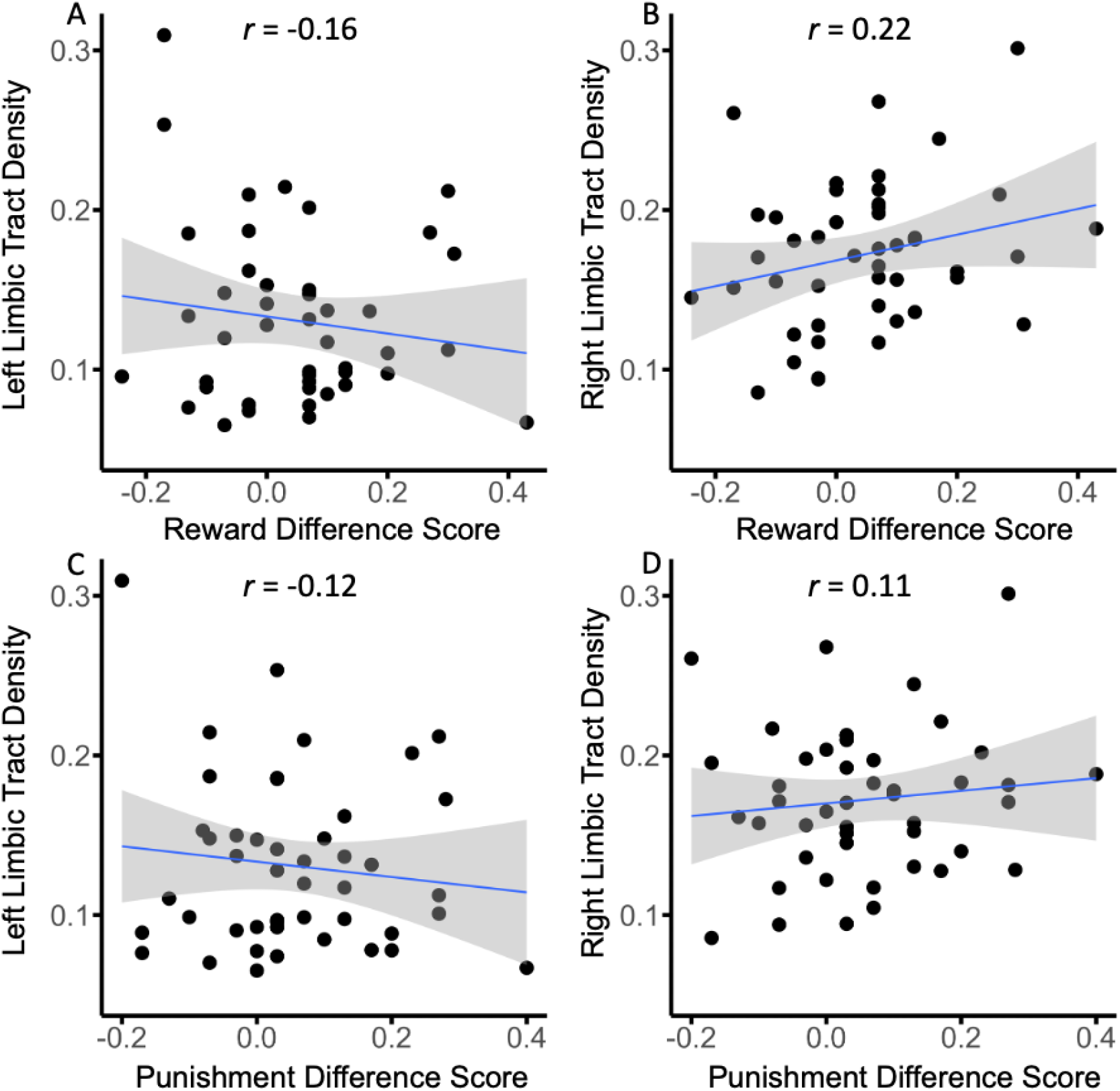
Scatterplots of reward (reward source memory hit rate minus control source memory hit rate) and punishment (punishment source memory hit rate minus control source memory hit rate) effects and limbic tract density. A) Left limbic tract density and reward effect B) Right limbic tract density and reward effect C) Left limbic tract density and punishment effect D) Right limbic tract density and punishment effect.

It is difficult to interpret null results when comparing two experimental effects. Therefore, we conducted Fisher r-to-z transformations to directly compare correlations between SN/VTA-hippocampus and SN/VTA-limbic pathways for both reward and punishment (Fisher, 1921). This analysis revealed a significant difference in the correlations for both reward (*Z* = 2.23, *p* < 0.05, two-tailed) and punishment (*Z* = 2.11, *p* < 0.05, two-tailed) in the left hemisphere.

No significant difference was found for reward (*Z* = 0.52, *N*.*S*., two-tailed) or punishment (*Z* = 1.11, *N*.*S*., two-tailed) in the right hemisphere. We also conducted analyses to test whether individual differences in memory prioritization were related to the other SN/VTA-striatum tracts (SN/VTA-executive, SN/VTA-motor). The measures were exploratory, and we expected no significant correlations to emerge. We computed partial correlations between the reward effect and SN/VTA-striatum tract density and the punishment effect and SN/VTA-striatum (SN/VTA-executive, SN/VTA-motor) tract density for each hemisphere (controlling for intracranial volume). The results revealed that neither reward nor punishment effects were correlated to SN/VTA-striatum tract density.

## Discussion

We used diffusion weighted MRI and probabilistic tractography to map the topology of the midbrain SN/VTA and to relate structural connectivity measures to individual differences in reward- and punishment-motivated memory encoding. Behaviorally, both reward and punishment effects were localized to associative memory (correct source recognition) and not found for item memory. These findings support previous findings that reward- and punishment-motivated encoding selectively enhances deeper, recollective memories (Elliott & Brewer, 2019; Gruber & Otten, 2010; Hennessee et al., 2017). Memories that rely on recollection afford access to rich associative and contextual information from the study episode, whereas memories that rely upon familiarity only afford access to information of the study items. Different neural regions have been shown to dissociate recollection and familiarity. Recollection relies more upon the hippocampus, whereas familiarity relies more upon the perirhinal cortex (Diana et al., 2007). These results fit with the “binding of items and contexts” model. This model posits that the perirhinal cortex processes item information whereas the parahippocampal cortex processes context information. Finally, the hippocampus binds item and context information to support episodic memories (Ranganath, 2010). Our findings that reward- and punishment-motivated memory encoding selectively enhances source memory may be indicative of enhanced hippocampal processing, thereby permitting more enriched encoding episodes.

The neural mechanisms underlying motivated memory remain opaque. It is hypothesized that interactions between the dopaminergic reward system (SN/VTA and NAc) and the hippocampus are critical. Previous neuroimaging studies in humans have found the involvement of the SN/VTA and the NAc in the prediction of both rewards and punishments (Carter et al., 2009), and in learning from these predictions via reinforcement learning (Schultz et al., 1997).

Previous fMRI studies in humans during motivated memory encoding tasks have shown increased activation in the medial temporal lobe, SN/VTA and NAc are predictive of better memory prioritization for reward-predicting stimuli (Adcock et al., 2006; Wittmann et al., 2005) and punishment-predicting stimuli (Shigemune et al., 2014). Individual differences in functional connectivity between the SN/VTA and hippocampus have also been shown to predict individual differences in motivated memory performance (Wolosin et al., 2012).

Interpretations of the SN/VTA, NAc, and hippocampal activation observed during motivated memory encoding vary. Some researchers suggest that canonical SN/VTA-ventral striatum reward prediction errors (RPEs) underlie the reward-driven gain in episodic memory performance (Calderon et al., 2021; De Loof et al., 2018; Jang et al., 2019; Rouhani et al., 2018). Other researchers have theorized that the hippocampus-VTA loop is critical for motivated memory encoding (Adcock et al., 2006). This latter hypothesis suggests that dopamine released into the hippocampus via midbrain projections increases hippocampal plasticity, thereby enhancing encoding and consolidation of information into long-term memory (Lisman et al., 2011). However, these studies have all focused on functional activation within these circuits. Our findings suggest that neuroanatomical differences across people determine the strength of this reward-related effect on memory. We found that individual differences in tract density of the SN/VTA-hippocampus were positively correlated with both reward and punishment effects on human episodic memory. Additionally, Fisher’s r-to-z transformation revealed that these correlations were specific to the SN/VTA-hippocampus tracts and not the SN/VTA-limbic tracts (in the left hemisphere). The results provide evidence that structural connectivity of dopaminergic midbrain-hippocampus projections underlie reward-motivated memory encoding, rather, or in addition to, canonical SN/VTA-limbic RPE signaling.

Although other researchers have theorized SN/VTA-limbic striatal RPE signaling can drive memory performance (Calderon et al., 2021), we believe this may be a separate neural system that can influence episodic memory, depending on the task demands. Studies that find traditional SN/VTA-limbic striatum RPEs support episodic memory encoding typically implement a reinforcement learning paradigm with either instrumental or Pavlovian learning via feedback (Jang et al., 2019; Rouhani et al., 2018). Memoranda in these tasks are almost always incidentally encoded (Ergo et al., 2020). Studies have also found that hippocampal-driven episodic memory can compete for processing resources and impair striatal feedback-based learning (Foerde et al., 2013; Wimmer et al., 2014). The motivated source memory task implemented here requires participants to use effortful memory encoding strategies, which could bias activity in favor of the hippocampal-VTA episodic memory. Future studies are needed to further dissociate the exact contributions of the SN/VTA reward system to effortful episodic encoding versus conditioned learning.

Further evidence that motivated-memory encoding may have different neural generators from standard SN/VTA-striatum RPEs comes from a recent electroencephalography study (Elliott et al., 2020). This study parametrically manipulated reward value during the encoding period of a recognition memory task. The study found that a P3 event-related potential during encoding scaled linearly with the value of the studied item. Furthermore, individual differences in this P3 component were positively correlated with individual differences in reward-motivated memory performance. However, when we analyzed the feedback related negativity (FRN) component during encoding (an event-related potential component theorized to reflect canonical RPE signals transmitted from the midbrain to the ACC; Sambrook & Goslin, 2014; San Martín, 2012; Walsh & Anderson, 2012) we failed to find an effect of value. The findings suggest that different neural generators may code different types of reward value depending on the task (motivated memory versus reinforcement learning).

Studies have shown that connectivity between frontal cortex and striatum is defined by functionally disparate circuits that subserve sensorimotor, executive, and limbic functions. These parallel circuits are topographically organized in a dorsal-ventral configuration, with each structural circuit maintaining the functional characteristics of the target cortical region. Midbrain dopamine neurons display similar functional heterogeneous circuits with their projections to the striatum. The midbrain-striatum projections have a spiraling medial/lateral and inverse ventral/dorsal topographical arrangement from the VTA and SN. The dorsomedial regions of the SN/VTA innervate the ventral striatum and the ventrolateral regions innervate the dorsal striatum (Fallon & Moore, 1978; Lynd-Balta & Haber, 1994; McRitchie et al., 1996). This spiraling architecture is hypothesized to allow affective limbic signals to influence executive and motor actions (Haber & Knutson, 2010; Haber, 2016). We used tract density measures to test if connections to the hippocampus and striatum from the SN/VTA regions display the same anatomy known from primate and rodent histology. The SN/VTA-striatum connections revealed an inverse dorsal-ventral and medial-lateral configuration. The dorsomedial SN/VTA showed greatest connectivity to the limbic striatum, with more ventrolateral regions of the SN/VTA showing highest connectivity to the dorsal striatum. Importantly, connections to the hippocampus also fell primarily within the VTA and medial SN region. A recent study employing distinct but convergent diffusion weighted neuroimaging methods also revealed a medial-lateral gradient of midbrain-striatum pathways (MacNiven et al., 2020). The current findings replicate and extend previous findings that the medial-lateral anatomy of heterogeneous midbrain pathways (striatum and hippocampus) can be delineated non-invasively in a neurotypical human population (MacNiven et al., 2020; Elliott et al., 2022).

Anatomically, the hippocampus receives direct dopaminergic projections from the VTA (Bethus et al., 2010; Gasbarri et al., 1997; Luo et al., 2011; Zubair et al., 2021). To our knowledge, we are the first group to use probabilistic tractography to quantify SN/VTA-hippocampus tracts in a group of participants. Although tractography does not have the ability to resolve afferent and efferent connections, the hippocampus has no known direct efferent connections to the SN/VTA. Additionally, our results found that the hippocampus showed greatest connectivity with the dorsomedial SN/VTA, corroborating rodent and primate findings (Lisman & Grace, 2005, Zubair et al., 2021). We believe this is further evidence that the tracts quantified here do, in fact, represent dopaminergic VTA-hippocampus projections. The tractography results extend what is known from the animal literature to a neurotypical human population.

We found no significant correlations between reward and punishment effects and SN/VTA-striatum connectivity. Although previous fMRI studies implementing motivated-memory tasks have found limbic striatum (NAc) activation in addition to SN/VTA and hippocampus activation, this may not reflect the SN/VTA-limbic pathways quantified here. The hippocampus-VTA loop consists of two primary pathways: (1) the upward arc (VTA-hippocampus) that we quantified in this study and (2) the downward arc (hippocampus-NAc-ventral pallidum-VTA). If the hippocampus-VTA loop is requisite for motivated (reward and punishment) effects on episodic memory, it is possible that the NAc activation observed in previous studies is representative of activation along the downward arc (hippocampus-NAc connections), and not direct SN/VTA-NAc connectivity associated with RPEs, as often interpreted. Therefore, the NAc activation observed in other motivated memory paradigms could be epi-phenomenal, resulting from hippocampal signaling.

Memory is critical for goal-directed behavior, but human memory has a finite capacity. The excess of sensory information humans encounter daily requires that we selectively prioritize and encode important or rewarding information at the expense of less important information, allowing our memory systems to be adaptive. It is hypothesized that interactions between the dopaminergic reward system and hippocampus underlie memory selectivity. We found that both reward- and punishment-motivated memory encoding preferentially enhance associative memory and not item memory. Additionally, tract density of the midbrain-hippocampus connections was predictive of individual differences in motivated memory prioritization for both rewards and punishments. These results provide compelling evidence to support theories of SN/VTA heterogeneity and the involvement of the dopaminergic reward system in motivated memory encoding.

## Acknowledgements

This work was supported by an NSF GRFP (B.L.E.), NSF grants 1634179 and 2031708 (S.M.M.), and NIH grants R01 MH091068 and R01 AA027381 (S.M.M.).

