## Supplemental Material for "Midbrain-Hippocampus Structural Connectivity Selectively Predicts Motivated Memory Encoding" (from another version of this preprint: version 1, 2022-05-19)

**Supplemental Materials**

**SN/VTA-HPC Correlations**

For completeness, we replicated our correlation analyses between SN/VTA-hippocampus tract strength measures and individual differences in reward- and punishment-motivated memory encoding, including the outlier case that we removed. We computed partial correlations for the reward effect and SN/VTA-hippocampus tract integrity and the punishment effect and SN/VTA-hippocampus tract integrity for each hemisphere (controlling for intracranial volume). The results revealed that both the reward (*r*(43) = .409, *p* < .01) and punishment (*r*(43) = .426, *p* < .01) memory effects were positively correlated with the left SN/VTA-hippocampus tracts. Both effects did not reach statistical significance for the right SN/VTA-hippocampus tracts (*r*(43) = .196, N.S., *r*(43) = .191, N.S., Supplemental Figure 2).

**SN/VTA-Striatum Correlations**

Exploratory correlations were conducted between each SN/VTA-striatum tract and our individual difference measures for reward- and punishment-motivated memory encoding. We computed partial correlations for the reward effect and each SN/VT-striatum tract and the punishment effect and each SN/VT-striatum tract for each hemisphere (controlling for intracranial volume; Supplemental Figure 3). The results revealed that neither reward nor punishment effects were correlated to SN/VTA-striatum tract integrity (Supplemental Table 1).

We repeated these correlations with the outlier subject included (Supplemental Figure 4). The results revealed that neither reward nor punishment effects were correlated to SN/VTA- striatum tract integrity (Supplemental Table 2).

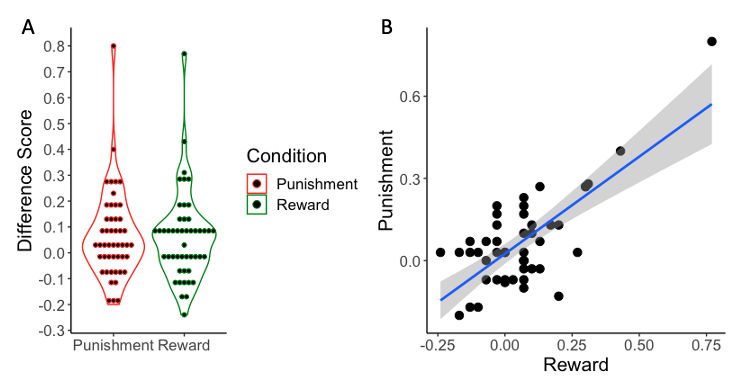

**Figure 1: Reward and punishment effects (difference scores) with outlier case included**. A) Violin plot B) Scatterplot.

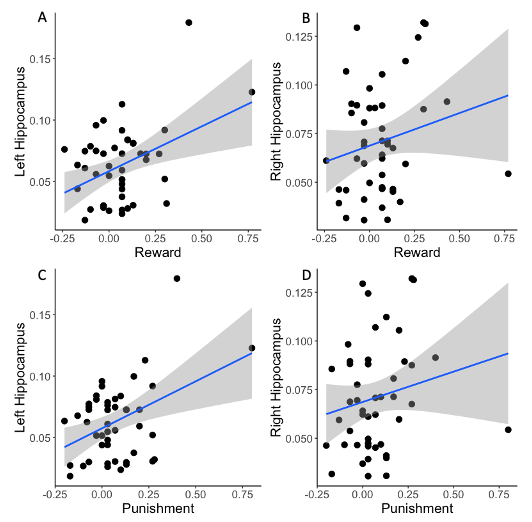

**Figure 2: Scatterplots of reward and punishment effects and hippocampus tract integrity with outlier included.** A) Left hippocampus tract integrity and reward effect B) Right hippocampus tract integrity and reward effect C) Left hippocampus tract integrity and punishment effect D) Right hippocampus tract integrity and punishment effect.

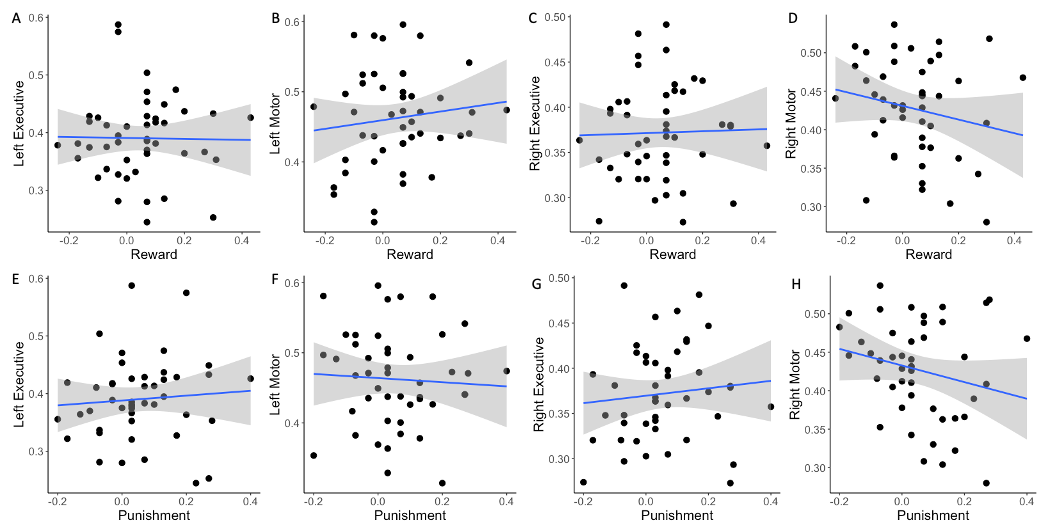

**Figure 3: Scatterplots of reward and punishment effects and striatum tract integrity.** A) Left executive tract integrity and reward effect B) Left motor tract integrity and reward effect C) Right executive tract integrity and reward effect D) Right motor tract integrity and reward effect E) Left executive tract integrity and punishment effect F) Left motor tract integrity and punishment effect G) Right executive tract integrity and punishment effect H) Right motor tract integrity and punishment effect

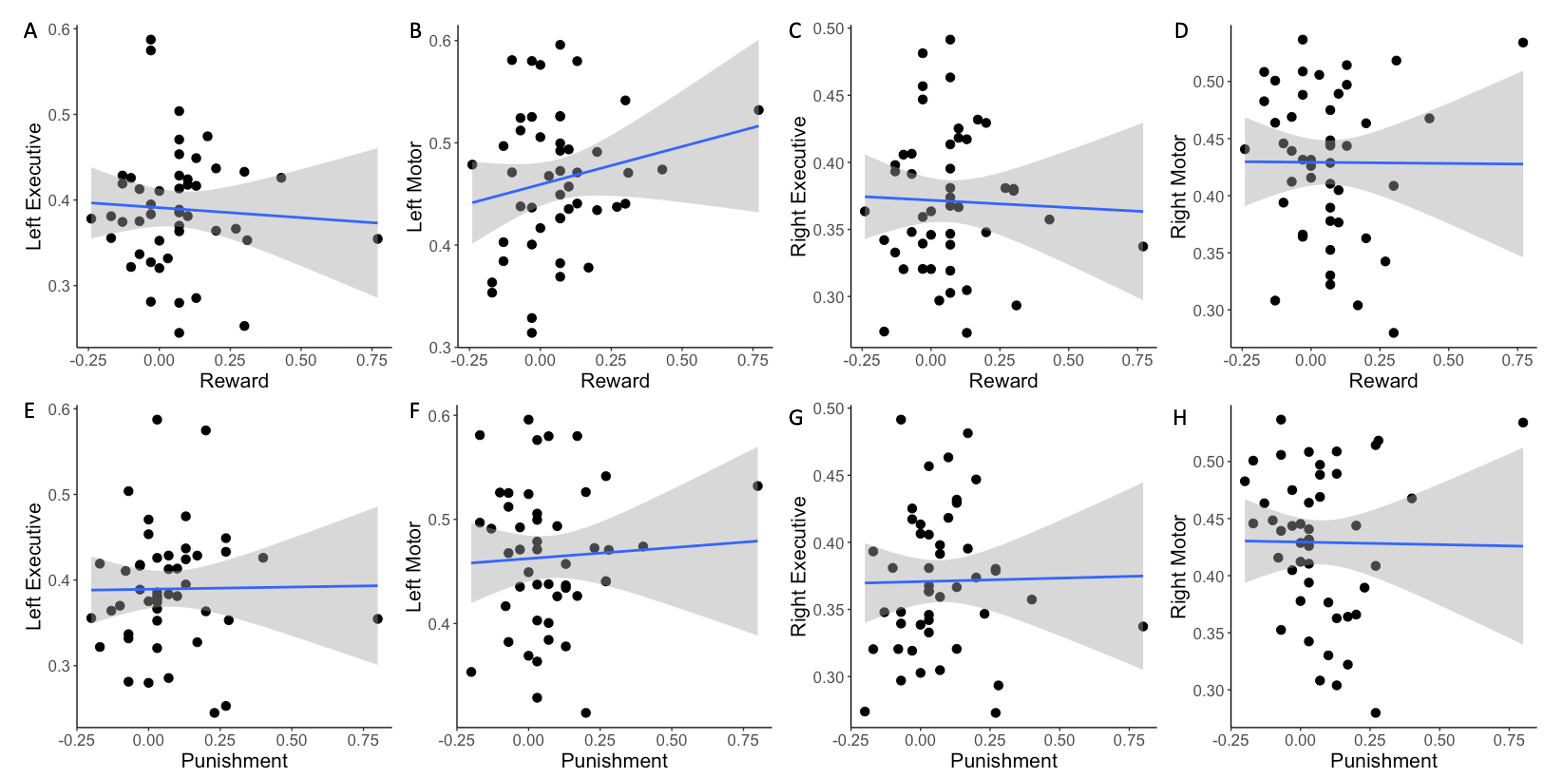

**Figure 4: Scatterplots of reward and punishment effects and striatum tract integrity (outlier included).** A) Left executive tract integrity and reward effect B) Left motor tract integrity and reward effect C) Right executive tract integrity and reward effect D) Right motor tract integrity and reward effect E) Left executive tract integrity and punishment effect F) Left motor tract integrity and punishment effect G) Right executive tract integrity and punishment effect H) Right motor tract integrity and punishment effect

**Table 1. Partial correlations between each SN/VTA-Striatum tract and reward and punishment effects (outlier removed).**

| Variables | | | Reward | Punishment | Left Executive | Left Motor | Right Executive | Right Motor |
| --- | --- | --- | --- | --- | --- | --- | --- | --- |
|  | Reward | Correlation | 1.000 | .550 | -.003 | .117 | .042 | -.200 |
|  |  | Significance (2-tailed) | . | <.001 | .983 | .450 | .788 | .192 |
|  |  | df | 0 | 42 | 42 | 42 | 42 | 42 |
|  | Punishment | Correlation | .550 | 1.000 | .080 | -.058 | .098 | -.219 |
|  |  | Significance (2-tailed) | <.001 | . | .604 | .710 | .527 | .154 |
|  |  | df | 42 | 0 | 42 | 42 | 42 | 42 |
|  | Left Executive | Correlation | -.003 | .080 | 1.000 | -.728 | .501 | -.184 |
|  |  | Significance (2-tailed) | .983 | .604 | . | <.001 | <.001 | .232 |
|  |  | df | 42 | 42 | 0 | 42 | 42 | 42 |
|  | Left Motor | Correlation | .117 | -.058 | -.728 | 1.000 | -.179 | .143 |
|  |  | Significance (2-tailed) | .450 | .710 | <.001 | . | .244 | .355 |
|  |  | df | 42 | 42 | 42 | 0 | 42 | 42 |
|  | Right Executive | Correlation | .042 | .098 | .501 | -.179 | 1.000 | -.612 |
|  |  | Significance (2-tailed) | .788 | .527 | <.001 | .244 | . | <.001 |
|  |  | df | 42 | 42 | 42 | 42 | 0 | 42 |
|  | Right Motor | Correlation | -.200 | -.219 | -.184 | .143 | -.612 | 1.000 |
|  |  | Significance (2-tailed) | .192 | .154 | .232 | .355 | <.001 | . |
|  |  | df | 42 | 42 | 42 | 42 | 42 | 0 |

**Table 2. Partial correlations between each SN/VTA-Striatum tract and reward and punishment effects (outlier included).**

| Variables | | | Reward | Punishment | Left Executive | Left Motor | Right Executive | Right Motor |
| --- | --- | --- | --- | --- | --- | --- | --- | --- |
|  | Reward | Correlation | 1.000 | .725 | -.047 | .181 | -.024 | -.008 |
|  |  | Significance (2-tailed) | . | <.001 | .758 | .233 | .876 | .958 |
|  |  | df | 0 | 43 | 43 | 43 | 43 | 43 |
|  | Punishment | Correlation | .725 | 1.000 | .014 | .051 | .014 | -.006 |
|  |  | Significance (2-tailed) | <.001 | . | .927 | .738 | .928 | .966 |
|  |  | df | 43 | 0 | 43 | 43 | 43 | 43 |
|  | Left Executive | Correlation | -.047 | .014 | 1.000 | -.729 | .504 | -.196 |
|  |  | Significance (2-tailed) | .758 | .927 | . | <.001 | <.001 | .197 |
|  |  | df | 43 | 43 | 0 | 43 | 43 | 43 |
|  | Left Motor | Correlation | .181 | .051 | -.729 | 1.000 | -.190 | .173 |
|  |  | Significance (2-tailed) | .233 | .738 | <.001 | . | .210 | .256 |
|  |  | df | 43 | 43 | 43 | 0 | 43 | 43 |
|  | Right Executive | Correlation | -.024 | .014 | .504 | -.190 | 1.000 | -.614 |
|  |  | Significance (2-tailed) | .876 | .928 | <.001 | .210 | . | <.001 |
|  |  | df | 43 | 43 | 43 | 43 | 0 | 43 |
|  | Right Motor | Correlation | -.008 | -.006 | -.196 | .173 | -.614 | 1.000 |
|  |  | Significance (2-tailed) | .958 | .966 | .197 | .256 | <.001 | . |
|  |  | df | 43 | 43 | 43 | 43 | 43 | 0 |
